# Collateral Responses to Classical Cytotoxic Chemotherapies are Heterogeneous and Sensitivities are Sparse

**DOI:** 10.1101/2021.04.15.440042

**Authors:** Simona Dalin, Douglas A. Lauffenburger, Michael T. Hemann

## Abstract

Chemotherapy resistance is a major obstacle to curing cancer patients. Combination drug regimens have shown promise as a method to overcome resistance; however, to date only some cancers have been cured with this method. Collateral sensitivity – the phenomenon whereby resistance to one drug is co-occurrent with sensitivity to a second drug – has been gaining traction as a promising new concept to guide rational design of combination regimens. Here we survey collateral responses to acquisition of resistance to four classical chemotherapy agents. Although collateral sensitivities have been documented for antibiotics and targeted cancer therapies, we did not observe collateral sensitivities to any of the cytotoxic agents we studied. Interestingly, we did observe heterogeneity in the phenotypic response to acquisition of resistance to each drug, suggesting the existence of multiple different states of resistance for each drug. Surprisingly, this phenotypic heterogeneity was unrelated to transcriptomic heterogeneity in the resistant cell lines. These features of phenotypic and transcriptomic heterogeneity must be taken into account in future studies of treated tumor subclones and in design of chemotherapy combinations.

## Introduction

Since the 1950s, clinicians have combated intrinsic chemotherapy resistance by treating patients with combinations of chemotherapies that have non-overlapping toxicities thereby allowing for higher doses. However, this method has resulted in an overall 5-year survival rate across all cancers of only 68%^1^. The antibiotic resistance field is facing a similar problem – rates of antibiotic resistance doubled between 2002 and 2014, and there are 23,000 deaths in the US each year due to antibiotic-resistant infections, according to the Centers for Disease Control and Prevention^2^. In response, researchers in the antibiotic resistance field have studied cases where resistance to one antibiotic causes, or is co-occurrent with, sensitivity to a second antibiotic – termed collateral sensitivity^3^. In theory, using collaterally sensitive drug pairs in combination or sequentially could select against the emergence of resistant microbes and thus reduce or prevent the emergence of antibiotic resistance^4^.

Recently, the concept of collateral sensitivity has begun to be explored in the context of cancer chemotherapy as well. For example, in 2016, Zhao *et al* found that in Philadelphia chromosome-positive acute lymphoblastic leukemia (ALL), resistance to dasatinib, a BCR-ABL1 inhibitor, can cause collateral sensitivity to crizotinib, foretinib, vandetanib, and cabozantinib - cMET and/or VEGFR inhibitors^5^. The following year, Andrew Dhawan and colleagues characterized collateral sensitivities to several tyrosine kinase inhibitors (TKIs) in ALK-positive non-small cell lung cancer (NSCLC). They found that cell lines resistant to first-line TKIs are often sensitized to non-TKIs including the classical chemotherapies etoposide and pemetrexed^6^. Finally, in 2020 a study from the same group showed that resistance to combinations of classical chemotherapies in Ewing’s sarcoma causes resistance to the drugs in the combinations, as well as to actinomycin D, and potentially sensitivity to SP-2509, a lysine-specific demethylase 1 inhibitor^7^.

While the growing knowledge of collateral sensitivities to targeted therapies is encouraging, most patients today are treated with at least one classical chemotherapy. A close review of the literature reveals several examples of likely collateral sensitivities between classical therapies. Vinblastine and paclitaxel have opposing mechanisms of action – the former destabilizes microtubules, and the latter stabilizes microtubules. They also have opposing mechanisms of resistance, vinblastine resistance can result from stabilizing mutations in α- and β-tubulin, while paclitaxel resistance can result from destabilizing mutations in α- and β-tubulin. When these resistance mechanisms are present, vinblastine-resistant cell lines are sensitive to paclitaxel, and vice versa^8^. Additionally, there is experimental and clinical data suggesting that, at least in some cases, cisplatin resistance can cause sensitivity to paclitaxel, and vice versa^9–11^. The mechanism behind this effect remains unclear, however, this combination has been found to be effective to treat patients with ovarian, breast, lung, skin, and head and neck tumors^12^.

While there are individual examples of collateral sensitivities to classical chemotherapies, there has not been a comprehensive survey of collateral sensitivities to single-agents in this class of drug, as has been done for antibiotics and targeted therapies. Furthermore, the dearth of known mutations that predict resistance to classical chemotherapies motivated us to define a proxy of resistant state by using the collateral effect of resistance to classical chemotherapies on sensitivity to other agents. Here, we survey collateral responses to acquired resistance to several classical therapies in the Eμ-Myc; p19^ARF -/-^ cell line which is a murine model for human Burkitt’s lymphoma^13,14^. We chose to investigate classical therapies which are commonly used in the clinic, several of which are used to treat Burkitt’s lymphoma, and several of which have collateral sensitivities described in the literature^15^. The drugs we studied are doxorubicin, a topoisomerase II poison, vincristine, a microtubule destabilizing agent, paclitaxel, a microtubule stabilizing agent, and cisplatin, a DNA crosslinker^16–18^.

Using a 96-well plate-based drug pulse and stepwise dose increase protocol, we were able to create >15 evolutionary replicates of stable cell lines resistant to each of these four drugs. We created a protocol to measure the resulting lines’ sensitivities to a panel of 20 chemotherapies with high precision. This work revealed marked heterogeneity in collateral changes in drug sensitivity between cell lines evolved to be resistant to the same drug. Transcriptomic profiling of cell lines resistant to paclitaxel was unable to explain the phenotypic heterogeneity. This heterogeneity in resistance acquisition needs to be taken into account when designing classical and salvage combination regimens in order to optimize patient outcome. Similarly, potential discordance between phenotype and transcriptome must be considered when studying transcriptome as a proxy for functional biology.

## Results

### Optimized protocol to evolve chemotherapy-resistant cell lines in vitro

To study collateral effects of resistance to classical chemotherapies, we developed a protocol allowing us to evolve many cell lines to resistance to one therapy in parallel. In these experiments we used the murine Eμ-Myc; p19^ARF -/-^ cell line as a tractable model because it is chemo-sensitive, has a short doubling time of ~12 hours, and grows in suspension so it can be easily manipulated in small culture wells^14^.

To evolve resistance to classical chemotherapies, 10,000 cells were treated with a dose of drug expected to kill 85% of cells over the course of three days in 96-well plates. After three days, killing was assessed using PI staining via flow cytometry and drug was removed from cells. Cells were allowed to recover for four days in untreated media and wells were split as they became confluent. On day 4, the plate was split to leave approximately 10,000 cells in each well which were treated again with a dose expected to kill 85% of cells based on the killing achieved the previous week. Remaining cells from the split were frozen and saved at −80°C (Fig 1A).

**Fig. 1.**
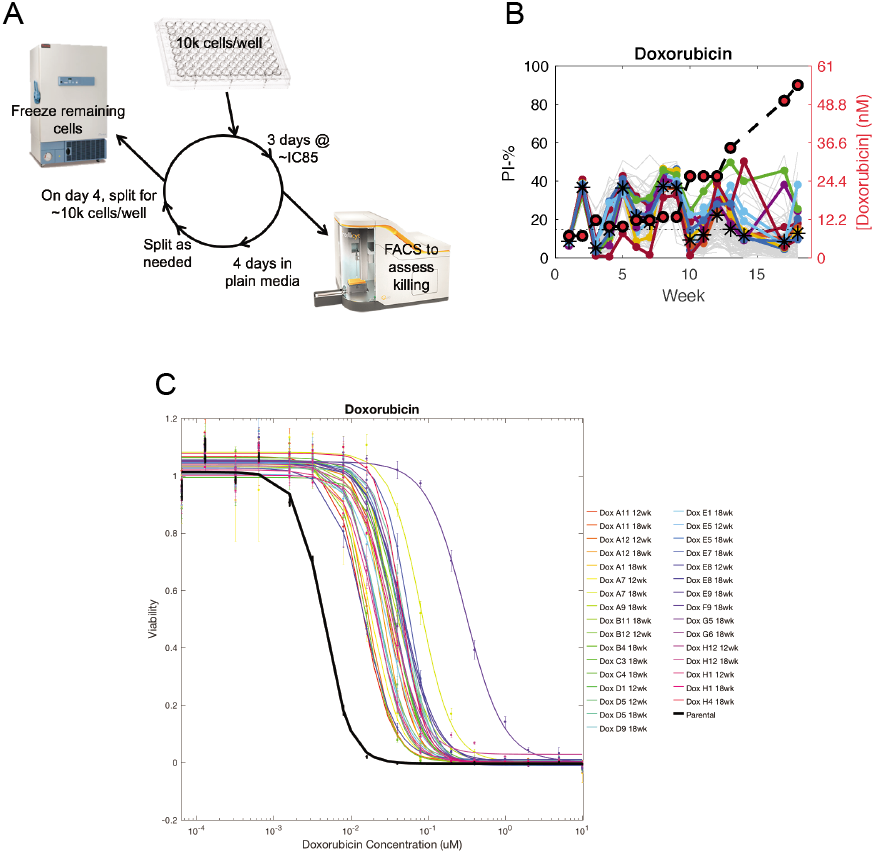
Protocol to create multiple evolutionary replicates of classical chemotherapy-resistant cell lines in parallel. **(A)** Schematic of protocol. In each cycle, 10,000 cells were seeded in each well of a 96-well plate and treated at a dose to kill 85% of cells after three days. Killing was assessed via FACS and cells were allowed to recover for four days. Cells were split, frozen, and the cycle re-started. **(B)** The fraction of live cells present in each well over each cycle of the resistance evolution experiment as assessed by PI staining (left y-axis). The dose of doxorubicin used each week is plotted with red circles on the right y-axis. Colored lines represent the most viable cell lines over the last five cycles of the experiment of the experiment. **(C)** Quantification of sensitivity to the selection drug after 12 or 18 cycles of selection.

As a pilot experiment, we used this protocol to create cell lines resistant to doxorubicin. Over the course of 18 weeks, cell lines were able to grow in increasing doses of doxorubicin and, by the end of the experiment, cell lines’ EC50 to doxorubicin increased by 9.7 ± 0.35 fold (Fig 1B,C).

### Screen of four doxorubicin-resistant cell lines against large drug library

To assess collateral changes in drug sensitivity, we screened four representative doxorubicin-resistant cell lines against an anti-cancer compound library containing 392 drugs. As expected, we observed collateral resistance to etoposide, another topoisomerase II inhibitor. We also observed a variety of other collateral changes in drug sensitivity, including collateral sensitivity to two HMG-CoA reductase inhibitors – fluvastatin and simvastatin (Fig 2).

**Fig. 2.**
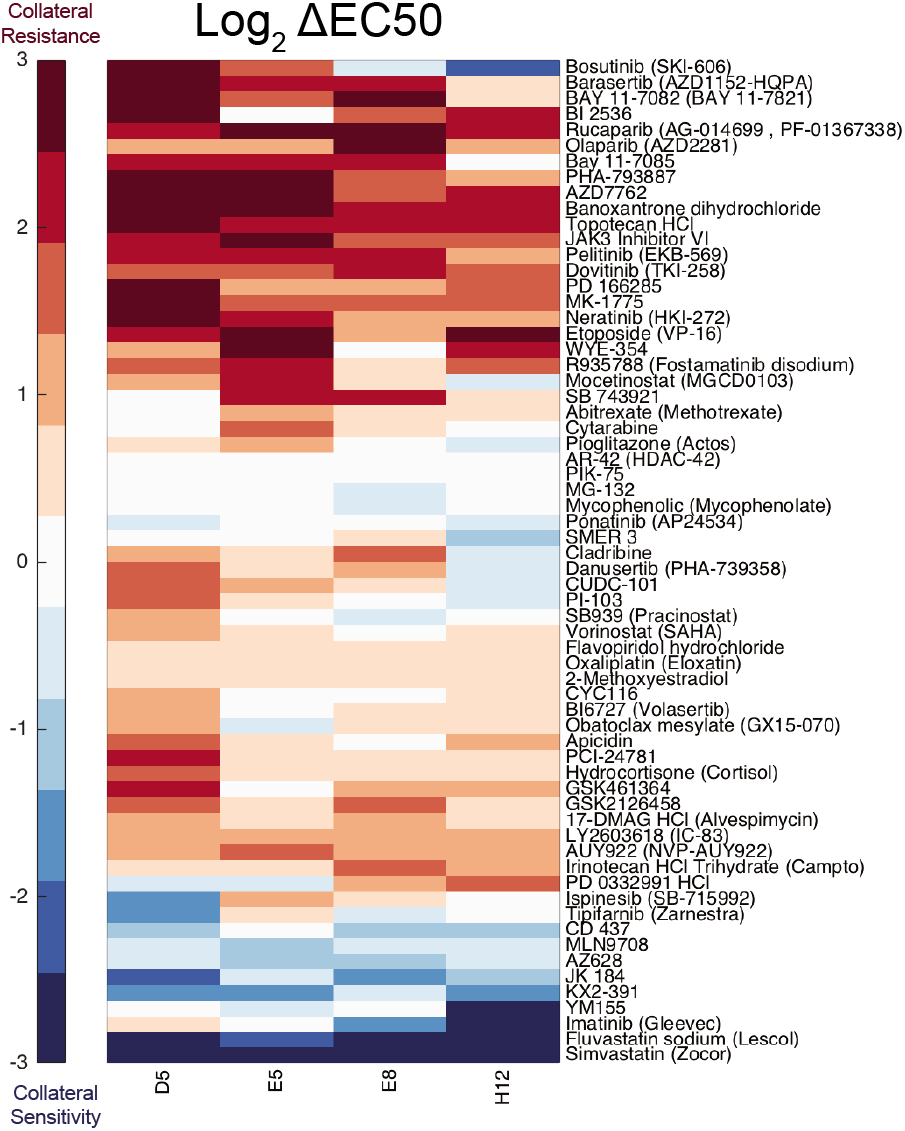
High-throughput drug screen to assess collateral responses to doxorubicin resistance. Cell lines screened are in columns, drugs with collateral effects are in rows. Colors represent the log_2_ fold change of the EC50 of each cell line for a drug relative to the parental cell line’s EC50 for that same drug. Red represents a case in which the resistant cell line also acquired resistance to a different drug – collateral resistance. Blue represents a case in which the resistant cell line acquired sensitivity to a different drug – collateral sensitivity.

### Generated cell lines resistant to four classical chemotherapies

We next used this method of creating resistant cell lines to generate cell lines resistant to vincristine, cisplatin, paclitaxel, as well as additional cell lines resistant to doxorubicin. We also generated control cell lines using DMSO in the place of drug. We were able to generate a total of 47 doxorubicin-resistant cell lines, 16 vincristine-resistant cell lines, 19 paclitaxel-resistant cell lines, 30 cisplatin-resistant cell lines, and 12 DMSO control cell lines. The fold change of EC50 to the selection drug was 206.5 ± 8.7 for doxorubicin-resistant cell lines, 10.82 ± 0.21 for vincristine-resistant cell lines, 5.2 ± 0.1 for paclitaxel-resistant cell lines, and 13.5 ± 0.3 for cisplatin-resistant cell lines (Fig 3A-D).

**Fig. 3.**
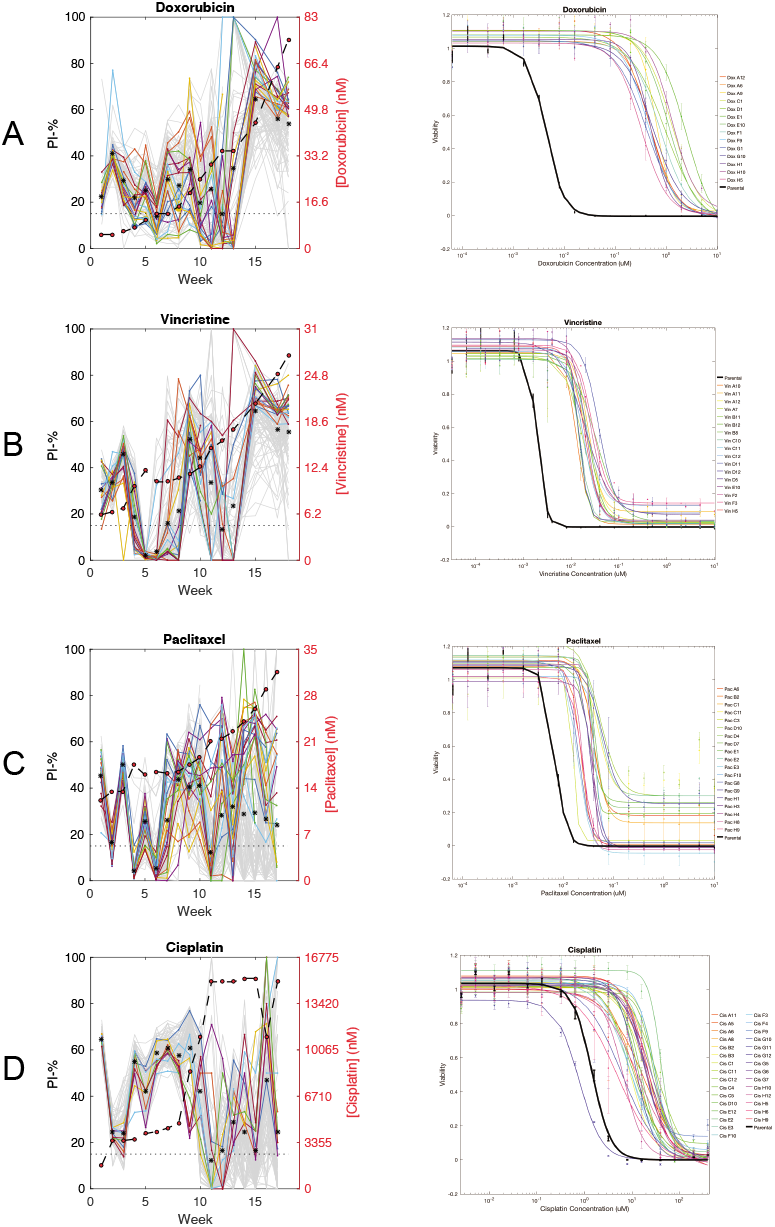
Viability of cell lines over each cycle of the resistance evolution experiment (left column) and quantification of sensitivity to the selection drug after 18 cycles (right column). In the left column, colored lines represent wells that were most viable over the last five weeks of the experiment. Each row represents acquired resistance to **(A)** doxorubicin, **(B)** vincristine, **(C)** paclitaxel, and **(D)** cisplatin.

### Developing a methodology to screen fewer drugs against many cell lines with high accuracy and precision

To determine collateral effects of resistance to doxorubicin, vincristine, paclitaxel, and cisplatin, we wanted to determine changes in sensitivity to chemotherapies. We developed a new dose response methodology to test a larger dose range and increase accuracy and precision of the drug sensitivity data relative to the screen we performed previously.

We reasoned that by choosing drugs that target a wide variety of biological processes, we may also gain insight into cellular processes changed over the course of acquiring resistance. For each of the chosen 20 drugs, we performed 25-fold 16-point serial dilutions in deep-well 96-well plates by hand. Each serial dilution was performed over four “quadrant plates”. We then used a Tecan liquid handler to transfer 20μl of that serial dilution to 384-well plates. 30μl of cells were plated on top of drug using a BioTek plate washer. After three days of incubation, we assessed viability with resazurin using a Tecan plate reader.

### Resistant cell lines exhibited extensive collateral resistance and limited collateral sensitivity

To study collateral changes in drug sensitivity resulting from classical chemotherapy resistance, we processed the raw dose response data into EC50 values for each drug and cell line. We then calculated the log_2_ fold-change in EC50 values for each cell line and drug pair relative to the parental cell line’s EC50 value for the same drug.

We observed extensive collateral changes in response to resistance to each of the four drugs we studied here (Fig 4A-D). Most collateral effects were in the form of resistance; however, we also observed collateral sensitivity to several drugs, most notably to the HMG-CoA inhibitors fluvastatin and simvastatin. Collateral sensitivity to these two drugs was observed in about half of all the resistant cell lines we studied regardless of the drug used to induce resistance. Upon further study, we noted that about half of the DMSO control cell lines also exhibited increased sensitivity to these two drugs, suggesting that this observation is an artifact of the culturing system we designed. The same appears to be true for the survivin inhibitor YM155.

**Fig. 4.**
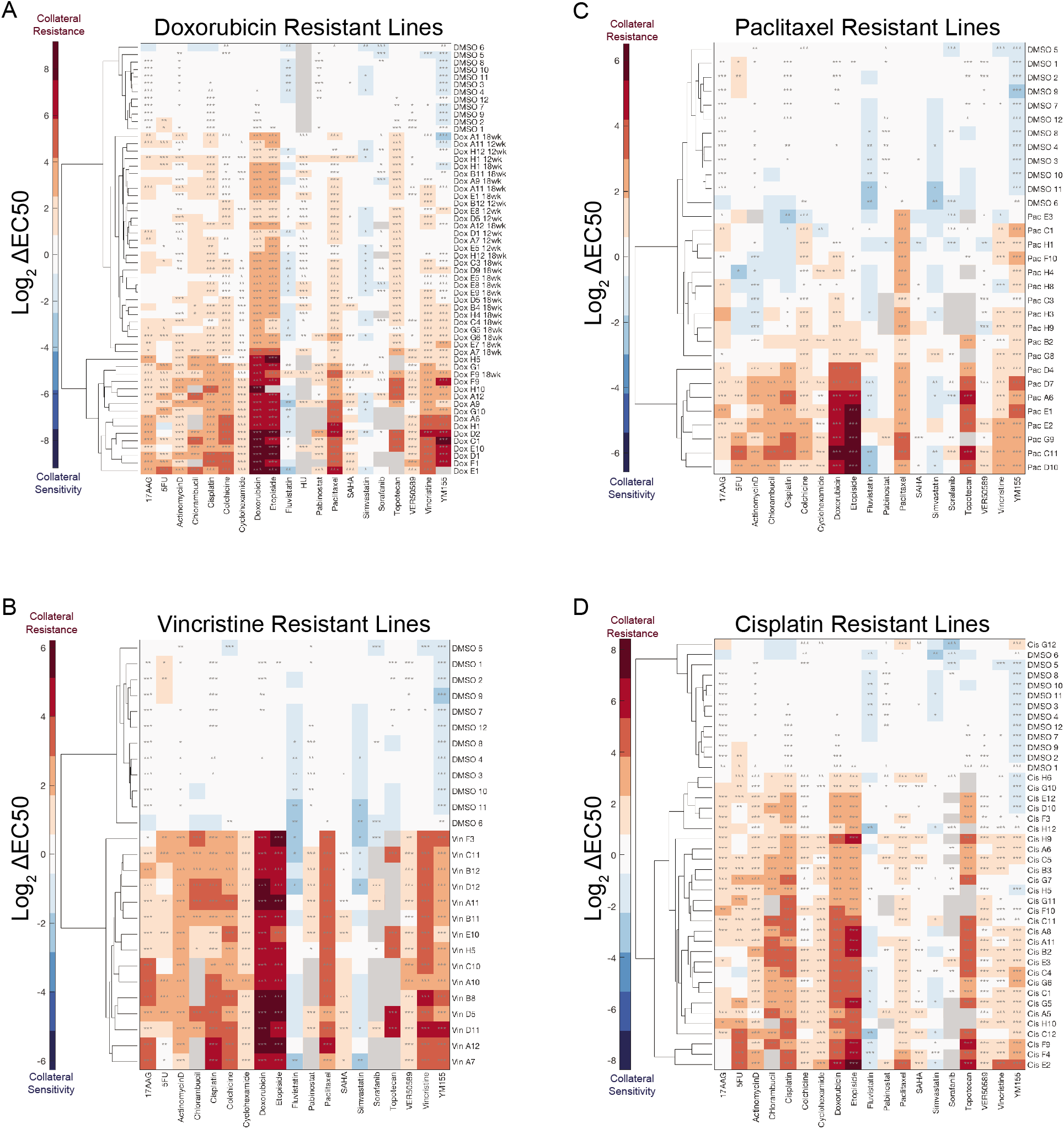
Collateral effects of acquiring resistance to **(A)** doxorubicin, **(B)** vincristine, **(C)** paclitaxel, and **(D)** cisplatin. In each plot, control cell lines cultured with DMSO instead of drug are shown at the top. Colors represent the log_2_ fold change of the EC50 of each cell line for a drug relative to the parental cell line’s EC50 for that same drug. Data represent the mean of three biological replicates. *, P ≤ 0.05, **, P ≤ 0.01, ***, P ≤ 0.001 (two-sample t-tests with Bonferroni correction).

### Collateral effects of classical chemotherapy resistance are heterogeneous

We observed notable heterogeneity in the collateral drug responses we measured. For example, the paclitaxel-resistant cell lines exhibited a wide range of sensitivities to Actinomycin D (Fig 4C). To formally assess this heterogeneity, we performed unsupervised hierarchical clustering on the log_2_ fold-changes in sensitivity to the drugs studied (Fig. 4A-D). In the case of doxorubicin and paclitaxel resistance, hierarchical clustering demarcates two clusters of resistant cell lines with distinct collateral drug sensitivity signatures (Fig 4A, C). This clustering methodology did not clearly distinguish any clusters of vincristine- or cisplatin-resistant cell lines. However, for each drug assayed, we are still able to observe a range of changes in EC50 values. This suggests that isogenic cells can respond to the same evolutionary pressure in heterogeneous ways, resulting in diverse phenotypes of the resulting resistant cells.

### Phenotype-based clusters of paclitaxel resistant cell lines are not associated with transcriptomic-based clusters of the same cell lines

To study the cause of the observed phenotypic clustering of resistant cell lines, we performed RNA-seq on the paclitaxel resistant cell lines, the DMSO control cell lines, and the parental cell line. Two of the paclitaxel resistant cell lines did not pass QC metrics so only 17 of the paclitaxel resistant lines were sequenced. PCA of the transcriptomic data from these cell lines clearly separated the DMSO control lines from the paclitaxel resistant lines, and also revealed two groups within the paclitaxel resistant lines (Fig 5A). PCA of the phenotypic drug sensitivity data from these cell lines recapitulated the differences between the DMSO control vs. paclitaxel resistant cell lines, as well as the two groups of paclitaxel resistant cell lines we previously observed with hierarchical clustering (Fig 5B). Surprisingly, the phenotypic-based and transcriptomic-based groups within the paclitaxel resistant lines are independent of each other (Fig 5A-B). Despite this, in transcriptomic space, cell lines within phenotypic-based groups are significantly closer to other cell lines within their group than to cell lines in the other group (Fig 5C-D).

**Fig. 5.**
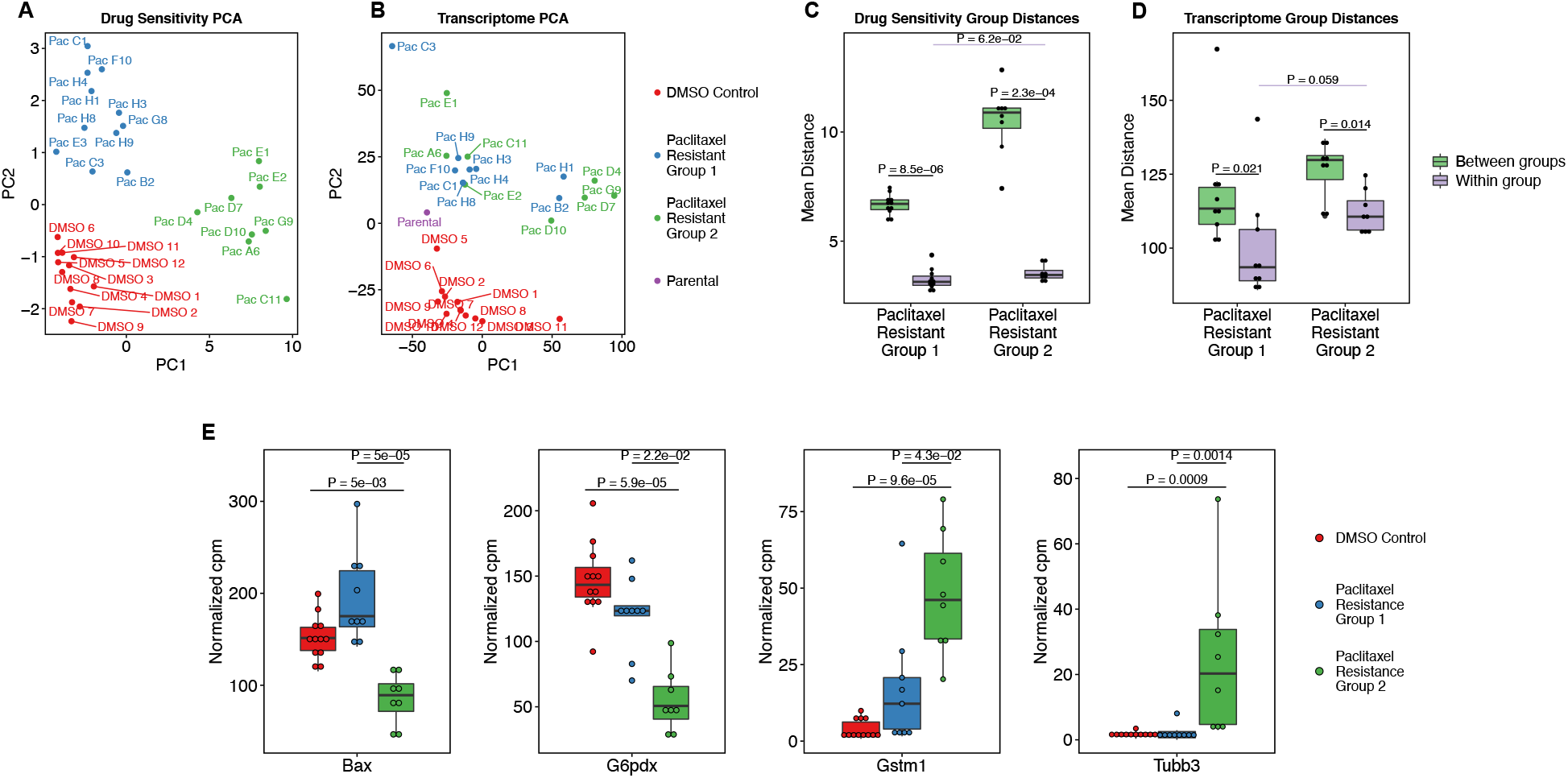
Phenotype-based groups of paclitaxel resistant cell lines. **(A)** PCA plots of log_2_ EC50 fold change values and **(B)** log of TMM-normalized counts-per-million expression values from RNA-seq for the paclitaxel resistant cell lines, DMSO control lines, and parental line. The blue and green annotated phenotypic groups are independent of the apparent transcriptome groups (Fisher’s Exact Test P = 0.33). **(C)** Mean Euclidian distance between cell lines within the phenotypebased paclitaxel resistance groups and **(D)** between the groups in drug resistance space, and in transcriptome space. FDR-corrected P-values are reported for two-sample Wilcoxon tests. **(E)** TMM-normalized counts for genes with a significant Kruskal-Wallis test and significant post-hoc Dunn’s test (Supplementary Fig S1B) are shown. FDR-corrected P-values are shown for post-hoc Dunn’s test of the indicated comparisons.

### Gene expression sheds light on different resistance mechanisms between phenotype-based groups of paclitaxel resistant cell lines

After observing two phenotype-based groups of paclitaxel resistant cell lines, we were interested in understanding transcriptome-based differences between these groups. However, these groups are interspersed with each other in transcriptome space, so global differential gene expression analysis revealed only two significantly expressed genes – Bst1 and Dglucy both show increased expression in Paclitaxel Resistant Group 1 (Figs 5, Supplementary Fig S1A & Supplementary Tables S4-6). Therefore, to assess additional potential differences in resistance mechanisms between these two groups, we compared expression levels of genes related to established mechanisms of paclitaxel resistance in these two groups as well as the DMSO control lines (Supplementary Table S1). Of the 26 genes with a significant difference in gene expression between these three groups of cell lines, only four, Bax, G6pdx, Gstm1, and Tubb3 showed a significant difference in expression between the two paclitaxel resistance groups (Fig 5E, Supplementary Fig S1B, & Supplementary Table S2).

## Discussion

We have established a method to reliably generate cell lines resistant to classical chemotherapies. We leveraged this technique to survey phenotypic responses collateral to acquiring classical chemotherapy resistance. Unexpectedly, we found that there was marked heterogeneity in collateral responses across cell lines exposed to the same evolutionary pressures. This heterogeneity suggests the existence of many possible cellular responses to the evolutionary pressure of chemotherapy. This hypothesis is supported by recent work showing that multiple mechanisms of resistance to a drug lead to stochastic collateral effects of resistance to that drug^19^.

The drugs we used in our screen have known resistance mechanisms that can be engineered in the lab such as topoisomerase II mutation or decreased expression causing doxorubicin resistance, and mutations in α- and β-tubulin^8,20,21^. However, after studying sensitivity to other compounds in our screen, in many cases, resistance did not appear to be based on these known mechanisms. For example, many doxorubicin and cisplatin resistant cell lines did not exhibit collateral resistance to 5-FU, a known P-gp drug efflux pump substrate^22^. Additionally, doxorubicin resistant cells were never collaterally sensitive to the topoisomerase I poison topotecan, as would be expected if resistance resulted from mutation or decreased expression of topoisomerase II^21^. Similarly, vincristine or paclitaxel resistance was not coincident with collateral sensitivity to the other drug, a result which would have been expected if resistance resulted from mutations in α- and β-tubulin. These results call into question, the clinical relevance of the known lab-based mechanisms of classical chemotherapy – if lab-based evolution of resistance does not result from the known mechanisms of resistance, does patient-based evolution of resistance result from those mechanisms? In fact, review of the literature reveals that the relevance of known resistance mechanisms is unclear in many cases, motivating further work to understand the nuances of clinically relevant collateral effects of chemotherapy resistance^23–25^.

The paucity of differentially expressed genes between the phenotype-based groups suggests that most of the alterations acquired during resistance evolution are passengers and unrelated to drug sensitivity. This is supported by our observations that the phenotypic-based and transcriptomic-based groups are not associated with each other. While the phenotypic-based groups do loosely cluster in transcriptomic space, it is clear that many of the alterations acquired are not associated with phenotypic response to other chemotherapies. This has important implications for study of tumor heterogeneity – transcriptomic profiling cannot be used as a proxy for phenotypic profiling because the relevant transcriptomic changes are obscured in a global analysis by unrelated changes caused by adaptation to the drug. In particular, certain approaches such as the connectivity map which seek to infer functional aspects of biology based on a transcriptional state may be misled by the myriad of unrelated variants^26^. By focusing on specific pathways known to be involved in paclitaxel resistance, we were able to show that one of the groups of paclitaxel resistant cell lines has changed state to become broadly chemotherapy resistant via changes in apoptotic priming, metabolism, drug efflux, and tubulin isoform expression. The cause of resistance in the other group of paclitaxel resistant cell lines remains unclear and will be investigated further in future work.

Further examination of the changes in drug sensitivity across our cell lines revealed increased sensitivity to two statins, which inhibit HMG-CoA reductase, in about half of the resistant cell lines. At first glance, this suggested an interesting avenue of rationally designing a combination regimen aimed at reducing chemotherapy resistance – adding a statin to a classical chemotherapy regimen may specifically target resistant cells. However, about half of the cell lines cultured in DMSO also exhibited increased sensitivity to these statins, suggesting that this phenotype is a result of the culturing system rather than a collateral effect of chemotherapy resistance. Recently there has been evidence in the literature that statins may be effective as cancer therapies; however, these results need to be carefully considered if culturing techniques can effect cell lines’ statin sensitivity^27–30^.

After removing artifact drugs for which the DMSO control cell lines exhibited changes in sensitivity, we observed collateral resistance in our resistant cell lines, but no collateral sensitivities. This is surprising, as collateral sensitivity has been widely documented in response to antibiotic resistance^31–34^, as well as to targeted chemotherapy resistance^6,35^. While the concept of collateral sensitivity has promise as a method to rationally design antibiotic or targeted therapy combination regimens, our result suggests that collateral sensitivity will be less useful for the rational design of classical chemotherapy combination regimens. In the clinic, poor response to 1^st^-line treatment with classical chemotherapy is predictive of failure of 2^nd^-line therapies as well. This observation predicts the result we observed but is complicated by the fact that most patients are treated with more than one agent. The fact that collateral sensitivity to classical chemotherapies is rare even in the single agent setting tested here underlines the reality that our understanding of the short- and long-term effects of these drugs is lacking. In contrast, resistance to targeted therapies is much better understood, and in fact targeted therapies induce collateral sensitivities. Perhaps the early use of classical chemotherapies in treatment is problematic and should instead be a last resort. Instead, to extend patient survival, frontline therapy should consist of combinations of targeted therapies which have more predictable resistance mechanisms and can result in treatable collateral sensitivities.

The sparsity or complete absence of collateral sensitivities to classical chemotherapies distinguishes the actions these drugs exert on cells from those exerted by antibiotics or targeted therapies. While antibiotics and targeted therapies are thought to interact relatively specifically with their targets, classical chemotherapies are thought to be more promiscuous and thus have a wider distribution of binding affinities for entities within the cell^36–38^. Our results lead us to hypothesize that collateral sensitivities result from strong evolutionary pressure on one or a small number of specific targets within the cell. In these cases, resistance can result from on-target mutations or target bypass, both of which can produce novel and exploitable sensitivities^35,39–41^. In contrast, the diffuse evolutionary pressure exerted by a compound with many binding partners would not be addressed by any single mutation or bypass mechanism as many cellular processes are affected. In this case, resistance may instead result from a more general state change, for example, to an anti-apoptotic state which would not be vulnerable to inhibition of any specific node within the cell. This paradigm leads to our observed result – few or no collateral sensitivities to classical chemotherapies, while others have observed collateral sensitivities to antibiotics and targeted therapies.

Our results, showing lack of collateral sensitivities to classical chemotherapy resistance, motivates further work to better understand mechanisms of action and resistance to these agents. Furthermore, efforts are needed to understand which combinations of classical chemotherapies work well together in order to rationally design novel combination regimens. This work is complicated by our observation of heterogeneity in phenotypic response to acquisition of classical chemotherapy resistance which is not explained by differences in gene expression. The discussion of heterogeneity in cancer has previously focused on inter-patient heterogeneity, heterogeneity between metastases within a patient, and intra-tumor heterogeneity. We have uncovered a new dimension of heterogeneity which needs to be further explored – heterogeneity in response and resistance to chemotherapy (Fig 6).

**Fig. 6.**
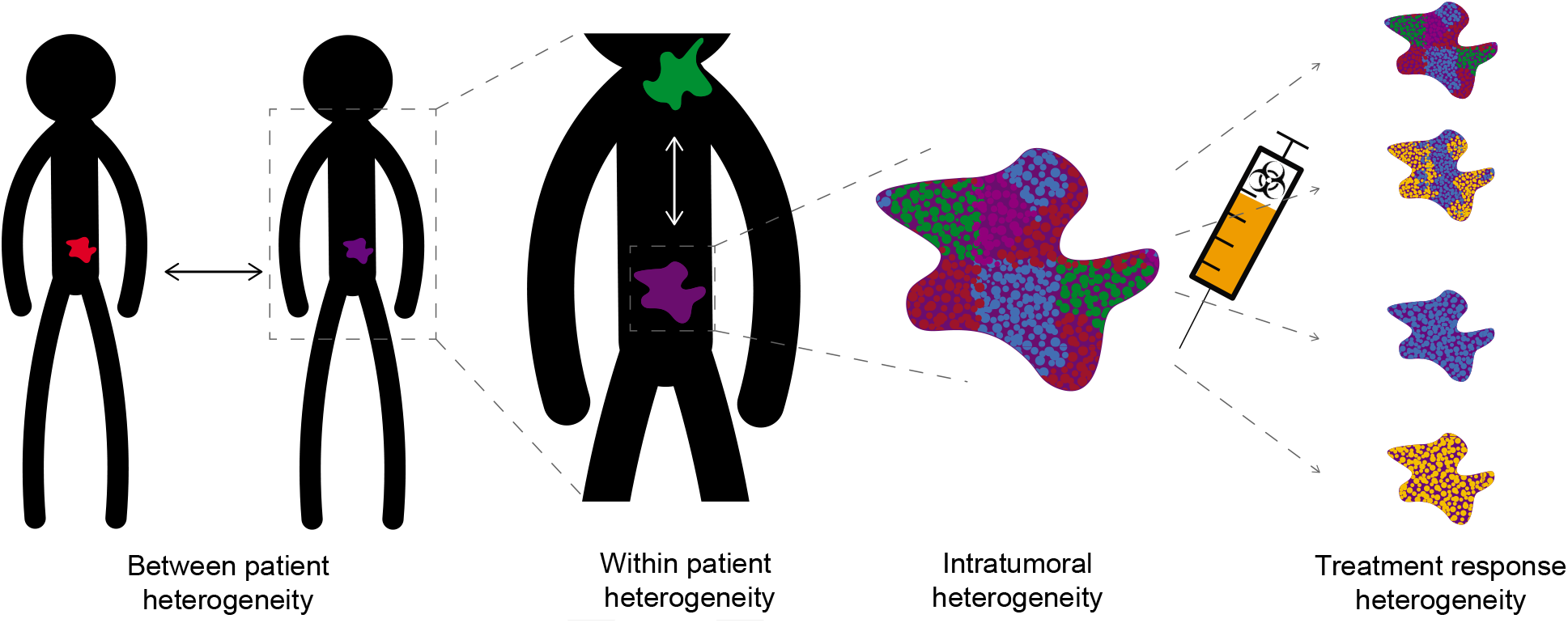
Schematic of a new dimension of tumor heterogeneity. Scientists and physicians have previously investigated heterogeneity between tumors in different patients, heterogeneity between different tumors in the same patient, and heterogeneity within the same tumor. Heterogeneity in tumor response to treatment also needs to be investigated.

## Materials and Methods

### Cell Culture and Chemicals

Murine Eμ-Myc; p19^ARF -/-^-B-cell lymphomas were cultured in B-cell medium *(45%* DMEM, *45%* IMDM, 10% FBS, supplemented with 2mM L-glutamine and 5μM β-mercaptoethanol). All cell lines were routinely tested for mycoplasma contamination using MycoAlert (Lonza). All drugs were obtained from LC Laboratories, Sigma-Aldrich, Calbiochem, or Tocris Biosciences.

### Creation of resistant cell lines

10,000 cells were seeded into each well of 96-well plates (Falcon), then treated with a dose of a chemotherapeutic to kill 85% of cells after three days. After treatment, viability was assessed via PI staining and FACS. Cells were spun out of drug and allowed to recover in drug-free media for four days. During recovery, cells were split as individual wells’ media color started to change. On day four, all wells were split to approximately 10,000 cells per well and the removed cells were frozen at −80°C in PCR plates. The cycle was repeated for 12-18 weeks.

### Cell Viability Assay

Cells were seeded in cell-culture treated 384-well plates (Falcon), then cells were treated as indicated. Cell viability was measured after 72 hours using resazurin sodium salt (Sigma). Resazurin was used at 0.008 mg/mL and plates were analyzed 6 hours after addition. Fluorescence was measured using the Tecan M200 Pro at an excitation of 550nm and emission of 600nm.

### Calculation of EC50s

The following analysis was performed in Matlab: median fluorescence of wells without cells was subtracted from all wells on the same plate. Viability was then normalized to cell growth by dividing all corrected fluorescence values by the median corrected fluorescence of wells containing untreated cells. Viability data was then fit to the following four-parameter hill curve:

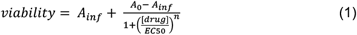

Where A_inf_ is the viability at high doses of the drug, and A_0_ is the viability with low or no drug. Sensitivity metrics were generated for each replicate individually, which were then used for further statistical analyses.

### RNA-sequencing and Differential Gene Expression Analysis

Total RNA was prepared from the paclitaxel resistant cell lines, the DMSO control cell lines, and the parental cell line using the RNeasy Mini Kit (Qiagen), per the manufacturer’s protocol, and included the on-column DNAse step (Qiagen). RNA samples were quantified and quality assessed using an Advanced Analytical Fragment Analyzer. The initial steps were performed on a Tecan EVO150.10ng of total RNA was used for library preparation. 3’DGE-custom primers 3V6NEXT-bmc#1-15 were added to a final concentration of 1 uM. (5’-/5Biosg/ACACTCTTTCCCTACACGACGCTCTTCCGATCT[BC_6_]N_10_T_30_VN-3’ where 5Biosg = 5’ biotin, [BC6] = 6bp barcode specific to each sample/well, N10 = Unique Molecular Identifiers, Integrated DNA technologies), to generate two subpools of 15 samples each.

After addition of the oligonucleotides, Maxima H Minus RT was added per manufacturer’s recommendations with Template-Switching oligo 5V6NEXT (10uM, [5V6NEXT: 5’-iCiGiCACACTCTTTCCCTACACGACGCrGrGrG-3’ where iC: iso-dC, iG: iso-dG, rG: RNA G]) followed by incubation at 42C for 90’ and inactivation at 80C for 10’.

Following the template switching reaction, cDNA from 15 wells containing unique well identifiers were pooled together and cleaned using RNA Ampure beads at 1.0X. cDNA was eluted with 17 ul of water followed by digestion with Exonuclease I at 37C for 30 minutes, and inactivated at 80C for 20 minutes.

Second strand synthesis and PCR amplification was done by adding the Advantage 2 Polymerase Mix (Clontech) and the SINGV6 primer (10 pmol, Integrated DNA Technologies 5’-/5Biosg/ACACTCTTTCCCTACACGACGC-3’) directly to the exonuclease reaction. 8 cycles of PCR were performed followed by clean up using regular SPRI beads at 0.6X, and eluted with 20ul of EB. Successful amplification of cDNA was confirmed using the Fragment Analyzer^42,43^.

Illumina libraries were then produced using standard Nextera tagmentation substituting P5NEXTPT5-bmc primer (25μM, Integrated DNA Technologies, (5’-AATGATACGGCGACCACCGAGATCTACACTCTTTCCCTACACGACGCTCTTCCG*A*T*C*T*-3’ where * = phosphorothioate bonds.) in place of the normal N500 primer.

Final libraries were cleaned using SPRI beads at 0.7X and quantified using the Fragment Analyzer and qPCR before being loaded for paired-end sequencing using the Illumina NextSeq500 in paired-end mode (26/50 nt reads).

Quality control was done with FastQC version 0.11.4^44^. Single-end reads were aligned to the GRCm38 mm10 assembly with gencode M25 annotation using STAR version 2.5.1b^45^. Gene counts from STAR were used to perform differential gene expression analysis with R version 3.6 using edgeR version 3.32.0^46,47^. Gene expression differences at the total gene level were considered significant at an FDR of < 0.01.

### Statistical Analysis

Statistics were performed using GraphPad Prism 5 (GraphPad Software Inc) and R version 3.6. All error bars represent standard error of the mean.

## Data Availability

The datasets generated during and/or analyzed during the current study are available in the GEO repository under accession number GSE166425.

## Acknowledgments

We would like to thank Boyang Zhao for many productive conversations, Yunpeng Liu for assistance with creating resistant lines, Rameen Beroukhim for transcriptome analysis suggestions, and Jackie Lees and Piyush Gupta for advising on this project. We thank the Koch Institute’s Robert A. Swanson (1969) Biotechnology Center for technical support, specifically Jaime H. Cheah and Christian K. Soule from the High Throughput Sciences Facility and Stuart Levine from the Genomics Core.

## Funding

This project was funded in part by the NIH (NCI U54-CA217377), the MIT Center for Precision Cancer Medicine, and by the Koch Institute Support (core) Grant P30-CA14051 from the National Cancer Institute. S.D. was supported by the David H. Koch Fellowship in Cancer Research. M.T.H acknowledges funding from the MIT Center for Precision Cancer Medicine and the Ludwig Center at MIT.

## Author contributions

S.D. conceived the project, designed and performed the experiments and analyses and wrote the manuscript. S.D., D.A.L., and M.T.H conceived the project and designed the experiments.

## Competing interests

The authors declare no competing interests.

## Data and materials availability

All data needed to evaluate the conclusions in the paper are present in the manuscript.

## Notes

**Additional Information:** Funding: This project was funded in part by the NIH (NCI U54-217377), the MIT Center for Precision Cancer Medicine, and by the Koch Institute Support (core) Grant P30-CA14051 from the National Cancer Institute. S.D. was supported by the David H. Koch Fellowship in Cancer Research. M.T.H acknowledges funding from the MIT Center for Precision Cancer Medicine and the Ludwig Center at MIT.

### Competing Interest Statement

The authors have declared no competing interest.

