## Supplemental Figure 1 for "Collateral Responses to Classical Cytotoxic Chemotherapies are Heterogeneous and Sensitivities are Sparse"

**Supplemental Figure S1**: Gene expression differences between phenotype-based paclitaxel resistant cell line groups

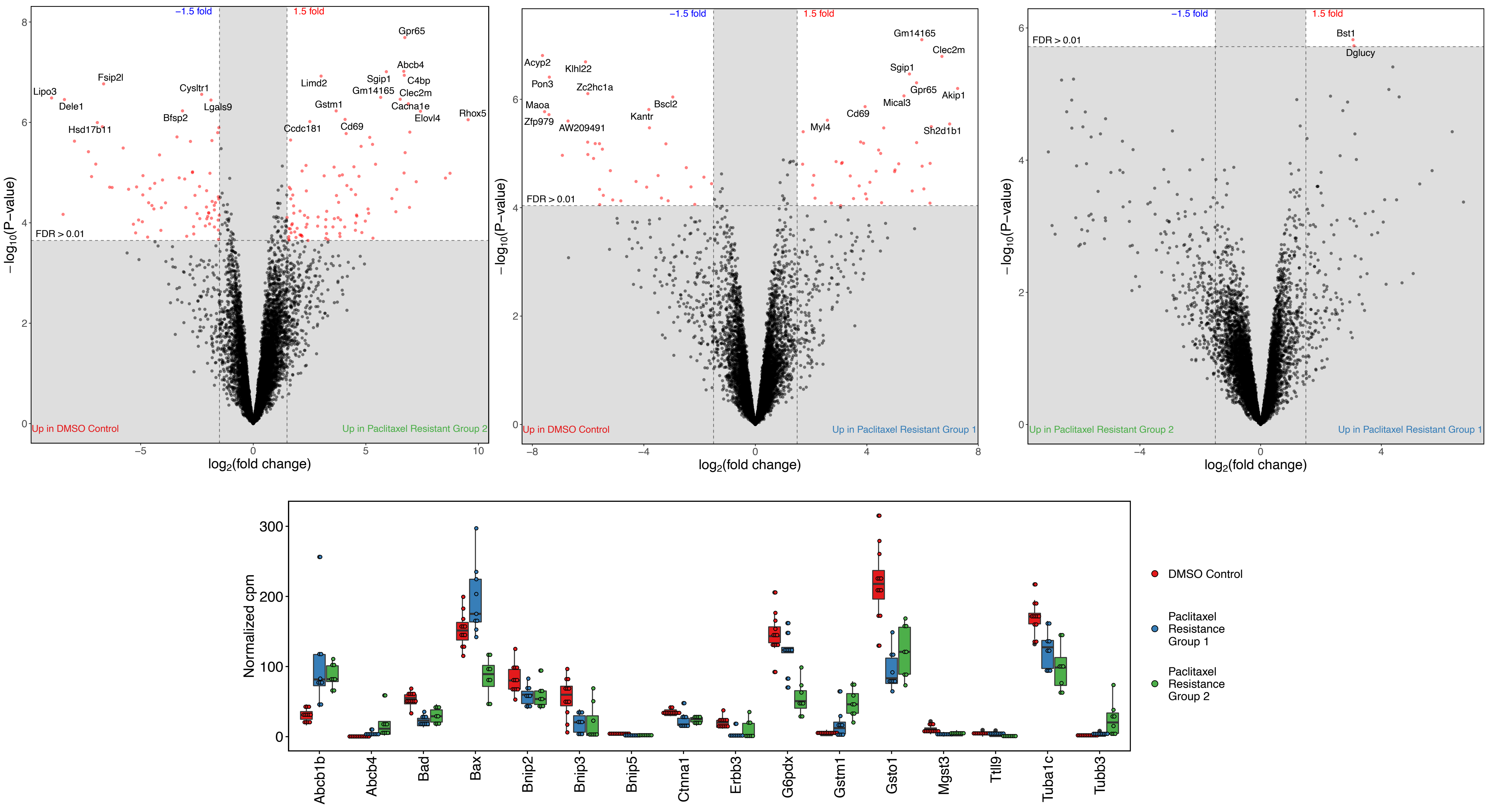
